# Comparison of a Computer Vision Model to a Human Observer in Detecting African Mammals in Camera Trap Images within a Safari Park

**DOI:** 10.1101/2025.06.02.656988

**Authors:** Naomi Davies Walsh, Carl Chalmers, Paul Fergus, Steven Longmore, Bridget Johnson, Serge Wich

## Abstract

Remote monitoring technologies are increasingly utilized in animal research for their capacity to enhance data collection efficiency. However, they present challenges, and as such researchers have resorted to utilizing deep learning to automatically classify acquired data therefore expediting the review process. While this practice is common in field studies it has been less adopted in zoo monitoring. In this paper we deploy the YOLOv10x model to monitor four species at Knowsley Safari in the UK: African lions (*Panthera leo*), Southern white rhino (*Ceratotherium simum simum*), Grevy’s zebra (*Equus grevyi*) and Olive baboons (*Papio anubis*). Camera trap images were processed and classified using the Conservation AI desktop application. The raw images were saved to facilitate the comparative analysis of the models’ predictions against the findings of human observed images. Processing time for both methods was compared using a subset of 3015 images with Conservation AI, reducing the time required to classify the images by 82% compared to a human analyst. Confusion matrix results showed high accuracy rates for all four species (>0.90). Analysis of count data showed significant differences in three species, where the human observer recorded more observations of each than Conservation AI (lion, rhino, baboon p<0.005). However, no significant difference was seen in zebra (p>0.05). A strong positive correlation in count data between both methodologies was seen in all species; baboon (rho=0.955, p<0.005), lion (rho = 0.969, p<0.005), rhino (rho=0.887, p<0.005) and zebra (rho=0.843, p<0.005). This study highlights the potential for these technologies as a monitoring system in zoos.

## 1. Introduction

### 1.1 Remote Monitoring of Species

To manage captive populations, provide exemplary care, and fulfil their role as conservation organisations, zoos need an evidence-based approach to husbandry that is specific to both species as a whole, as well as individual animals. To achieve this, widescale monitoring is vital (Congdon et al., 2022; Rose & Riley, 2021; Morrison & Novikova, 2022). Observation of animals can lead to pattern identification; presence or absence can help pinpoint welfare issues such as illness or stress, allowing zoos to take countermeasures to prevent suffering (Rose & Riley, 2021). Relevant member organisations and government licences also enforce monitoring and evidence-based reviews depending on location (Zoo Licencing Act, 1981). In person, direct observations require a lot of specially trained staff time (Diana et al, 2021). Many zoos use remote monitoring technologies, e.g., CCTV, wearable sensors, and microphones (Naidenov et al., 2024; Kalafut et al., 2020; Whitman & Miller, 2016) to mitigate the staff time required. Similarly, remote monitoring technologies are also popular for in-situ wildlife research (MacNulty et al., 2008; Trolliet et al., 2014; Wich et al., 2015; Kuster et al., 2020; Wich et al., 2023). In both in-situ and ex-situ settings, this is due to their ability to collect large volumes of important data, freeing up valuable time and reducing labour-intensive duties (Morrison & Novikova, 2022; Diana et al., 2021).

Camera traps, devices that use sensors (e.g., infrared) to trigger a recording of images or videos, are one such remote monitoring solution and have become a readily available and affordable way to monitor species in and exsitu. In situ, they are deployed in various habitats to monitor elusive species (Rovero & Zimmerman, 2016; Green et al., 2020; Newey et al., 2020; Morrison & Novikova, 2022). Camera traps are to conduct population estimates (Karanth et al., 1995; Rowcliffe et al., 2008; Harris et al., 2020), study behavioural ecology (Rowcliffe et al., 2016; McCarthy et al., 2019; Marion et al., 2022), investigate habitat use (Bowkett et al., 2008; Head et al., 2012; Godoy-Güinao et al., 2023), as well as surveying the impact of both positive human activities through conservation (Laneg et al., 2021; Littlewood et al., 2021), and negative human activities such as poaching or logging (Ramesh et al., 2017; Widness & Aronsen, 2018; Moore et al., 2021).

Ex-situ, camera traps are used to monitor breeding behaviours in zoos (Dedieu et al., 2023; Allan et al., 2018; Kachamakova et al., 2014), track movement of more elusive or nocturnal species (Baskir, 2021; Rose et al., 2018) and assess health and welfare to ensure effective husbandry (Smith et al., 2023). Camera traps can record the time and behaviour of an animal at a particular place and facilitate research without constant human presence. This is beneficial both in and ex-situ as manual observations often disturb animals and their habitats, leading to unintended impacts on the biodiversity of the area and/or natural behaviour of the species present (Rovero & Kays, 2021).

However, camera traps and other remote monitoring technologies are not without limitations (Harris et al., 2010; Sundaresan et al., 2011). They often produce large and difficult to manage volumes of data that need to be classified. The data collected can also include significant amounts of blank data where light, foliage, or shadows have triggered the camera (Meek et al., 2012; Burton et al, 2015). Most studies use multiple devices that are deployed for weeks at a time, and the subsequent large amount of data produced stored on either external or internal hard drives that are not easily accessible or sharable creating data silos (Ahumada et al., 2019). Typically, the data only provides a snapshot in time and is often centered around a single data source (Burton et al., 2015; Meek et al., 2015; Newey et al., 2015). The data is often hand-transcribed, leading to errors (Maydanchik, 2007), and drawing conclusions from these data can be time and resource heavy. This means that responses to current issues are reactive rather than proactive, often missing the opportunity for a successful intervention.

### 1.2 Integration of Machine Learning

Integrating computer vision is one solution to address the functional limitations of camera trapping. More studies are now utilizing deep learning (DL) approaches to automatically classify images and videos (Lamba et al., 2019; Vélez et al., 2023). This is perhaps the most effective methodology when considering both accuracy and reduction of required resources. Advances in DL architectures, combined with increased processing power such as GPUs, has meant that the technology has improved in picking out the constituent parts of an image, resulting in tremendous advances in fields such as facial recognition, medical image recognition and remote sensing (Rane et al., 2024).

Object detection, amongst other forms of data classification, has become increasingly popular in conservation research by assisting researchers in processing large volumes of data to answer pressing conservation and ecological questions (Green et al., 2020; Trnovszky et al., 2017). These applications include land degradation, biodiversity and habitat loss, trafficking of protected species, and tackling invasive alien species (Sisodia et al., 2023). When focussed on habitat and biodiversity loss, current developments in DL are facilitating the real time automation of animal detection and data analysis, to enable the automatic analysis of behaviours and the identification of abnormal events such as environmental disaster or illegal human activities such as poaching (Chalmers et al., 2019; Chalmers et al., 2021; Thompson, 2023; Fergus et al., 2024). This presents a unique opportunity to harness these technological advancements and build upon the way conservationists tackle long-standing issues relating to environmental protection and sustainable development (Vélez et al., 2023; Raihan, 2023). It is important to note that, as with all technology, object detection comes with its own set of limitations, one being a lack of training data that is representative of both the animal and the environment in which it operates, which leads to issues of misclassifications (Wearn et al., 2019). As such, creating a varied and representative dataset for the automatic species classification is important. However, this is challenging given the sparsity of particular species, often leading to class imbalances for underrepresented species. Zoos are able to contribute to data sets due to the ease of capturing images of a wide variety of species, especially in a multi-institutional effort.

Furthermore, object detection within an ex-situ setting could offer zoos a significant opportunity to move beyond manual image review and derive richer insights into animal movement patterns. Automating the identification of species within zoos makes it possible to gain insight into spatial use and interactions within enclosures, especially pertinent to mixed species exhibits (Fergus et al, 2024). These movement data could uncover subtle behavioral changes that might indicate stress, illness, or improvements in welfare following enclosure developments or environmental enrichment. Some zoos already use remote monitoring and artificial intelligence to answer such questions (Diana et al., 2021; Congdon et al., 2022); however, much of the published literature mainly discusses the potential benefits of real-time monitoring and automatic analysis without evidence of its widespread application. Expanding these technologies could provide an invaluable tool for zoos, combining efficiency with enhanced welfare monitoring.

### 1.4 Study Subjects and Aims

This paper will investigate the efficacy of automatic detection and classification in four commonly housed zoo species: African lions (*Panthera leo*), Southern white rhino (*Ceratotherium simum simum*), Grevy’s zebra (*Equus grevyi*), and olive baboons (*Papio anubis*). Although the study subjects have been chosen partly due to available data with the chosen zoo setting, three of the four species chosen are species of conservation concern, and therefore, additional long-term benefits to in-situ populations can also be drawn from this study (IUCN Red List, 2024). However, the primary purpose of this study is to assess the efficacy of the technology and discuss its ability to contribute to future monitoring advances in zoo environments.

Zoos provide a unique opportunity to study animal behaviour and welfare within a controlled environment. However, the effectiveness of such research relies heavily on precise and consistent monitoring, underscoring the necessity of advancing technological capabilities in zoos. This paper explores the potential of accelerated object detection technologies for enhancing research in zoos. We demonstrate the feasibility of species classification integrated with remote sensing technologies for efficient and continuous monitoring. Additionally, we discuss the opportunities of these technologies for baseline behavior and welfare analysis.

## 2. Materials and Methods

### 2.1 Data Collection and Classification

Data collection occurred at Knowsley Safari in Merseyside, UK, between 2021 and 2022, collected as part of a longer-term study. Seven cameras were used for this study: three in the African lion enclosure, two in the mixed species exhibit containing Southern white rhino, and one camera in each of the Grevy’s zebra and olive baboon enclosures. Cameras were installed outdoors, set up to focus on high-use areas; night quarters, access chutes, water troughs, and near houses (Figure 1). The cameras used for the study were ReoLink Go, 4G wireless cameras, fitted with a lithium battery (7800mAh Li9). ReoLink Solar Panels were fitted to recharge the battery, and a Vodafone SIM Card was inserted to provide data for real-time image transmission through the mobile network. The cameras captured images at a resolution of 1920 × 1072 pixels. The infrared (IR) sensor sensitivity was set to medium.

**Figure 1.**
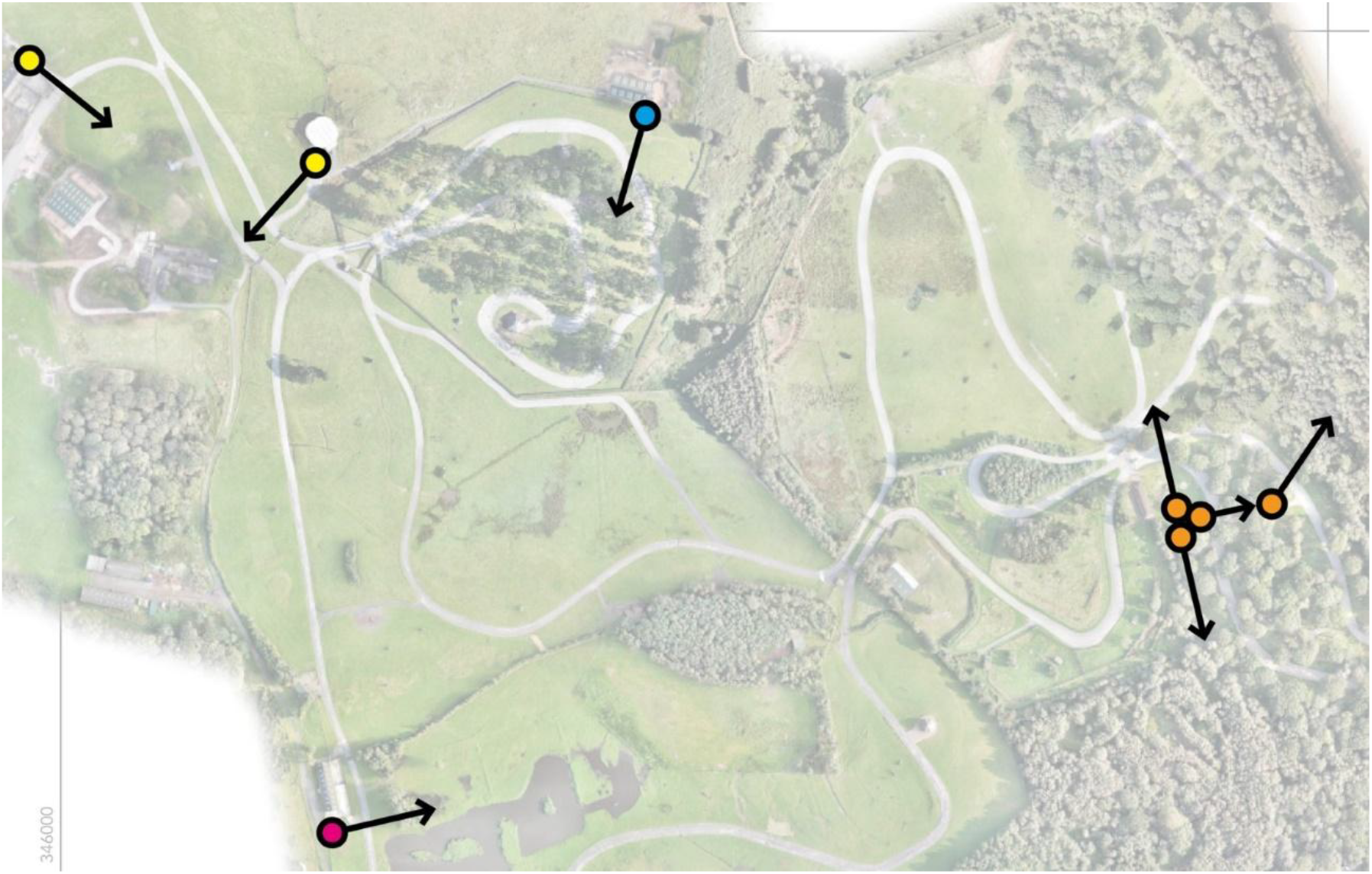
Google image view of the relevant enclosures at Knowsley Safari, with the camera positions and visual direction highlighted for lions (orange), rhino (yellow), baboons (blue), zebra (pink) (Google Maps, accessed 03.09.2024).

At the time of this study, Knowsley Safari housed six African lions, seven Southern White rhino, two Grevy’s zebra, and ∼215 Olive baboons kept in different sections of the safari drive (Figure 1). The lions were housed together in night quarters comprising of a house and outdoor paddock overnight and let out into a single species drive-through safari reserve daily during visitor opening hours. The rhino were housed in a single-species house overnight, then let out into two separate mixed-species exhibits with a range of large African mammals during visitor opening hours. The zebra were kept in a mixed-species house overnight and released into a single-species enclosure during visitor opening hours. Finally, the baboons had free access to two houses and were kept in a single species drive-through exhibit during the day and night. Due to housing differences, only lion and baboon images were collected during closed periods. The combined dataset used for this study contained 9644 images.

### 2.2 Model Selection and Inferencing

This study used a YOLOv10x model with a transfer learning approach for object detection and species classification. Details on hyperparameter selection and training were as detailed in Fergus et al. (2024). The trained model was hosted via the Conservation AI (CAI) website (www.conservationAI.co.uk), a publicly available site. This paper utilized the ‘Sub-Saharan’ model, which currently includes 32 species, humans, and cars. The model operates using a single-stage detection approach, simultaneously predicting both the location and identity of objects within an image. This contrasts with two-stage models such as Faster-RCNN, which separate region proposal from classification, often at the expense of processing speed. YOLOv10x contains 29.5 million trainable parameters, offering a strong balance between model complexity and computational efficiency. This allows for high-performance detection across large datasets without requiring specialized hardware (Wang et al., 2024). Its streamlined architecture and rapid inference capabilities make it particularly well-suited to real-time species monitoring where both speed and accuracy are essential, and multiple species may be present within the same image frame.

Inference was performed on a custom-built server featuring an Intel Xeon E5-1630v3 CPU, 256 GB RAM, and an NVIDIA Quadro RTX 8000 GPU. The model ran using the NVIDIA Triton Inference Server (v22.08) within a Docker container, operating on Windows Subsystem for Linux 2 (WSL2) (Savard et al., 2024). Images captured by camera traps in the field were transmitted via the Vodafone network to a centralized processing pipeline. Upon receipt by an SMTP server, images were routed through a web server which coordinated communication with a remote data center hosting the trained YOLOv10x model. Model inference was performed using NVIDIA Triton server with GPU acceleration via CUDA, enabling efficient object detection and species classification. If the prediction exceeded a 30% confidence rate, then the classified image and associated probability scores were loaded onto the CAI web platform under the assigned user’s account. If the probability score was below 30%, the image was classed as a blank and only uploaded if other species in the frame had a higher confidence rate. The resulting outputs, including bounding boxes and class labels, were then accessible through the CAI web platform for review.

### 2.3 Time comparison

For a subset of the data, 3015 images of baboons, a comparison was run assessing the time taken for the trained human observer to view and classify the number of baboons in each image, versus the time taken for them to upload and be classified on CAI. A stopwatch was running during each period of human data classification and was stopped for any disturbance. The stopwatch was also run during the upload period to CAI and stopped after the last file was processed.

### 2.4 Classification performance

Each image was viewed after classification on the CAI portal and assessed for accuracy of species detection by recording true positives (the target species was correctly classified) (TP), true negatives (another object or animal was classified correctly) (TN), false positives (another object or animal was incorrectly classified as the target species) (FP) and false negatives (the target species was incorrectly classified as another object or animal) (FN). These instances were recorded in a confusion matrix, and the following equations were used to assess the performance of the Sub-Saharan Africa model in accurately classifying each of the four included species:

1. Precision: Details the ability of the model to achieve a positive detection. Within this study, this demonstrates animals classified correctly, while accounting for any animals or objects that were incorrectly classified as the target species.

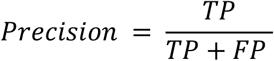
2. Recall: The proportion of positive detections that were correctly identified. Within this study, this uses the number of true positives against the number of false negatives to highlight the models’ ability to truly detect the target species.

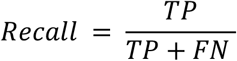
3. F1 Score: Provides a single score that balances both false positives and false negatives. A high F1 score in this study means the model can accurately classify and locate the target species within an image.

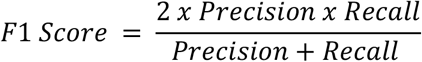
4. Accuracy: Summarizes the proportion of predictions that were correct (both positive and negative). It must however be noted that in imbalanced datasets, the accuracy of this metric can be misleading and should therefore be considered alongside others.

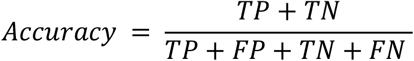

### 2.5 Count comparison: Conservation AI compared to a trained human observer

Once CAI had classified the images, a trained human observer viewed any images that included detections of each target species. CAI automatically filters out blank images, so these were not considered in this study. The number of detections in the image by CAI was recorded alongside the number of each target species the human observer could detect in the image (example: Figure 2).

**Figure 2.**
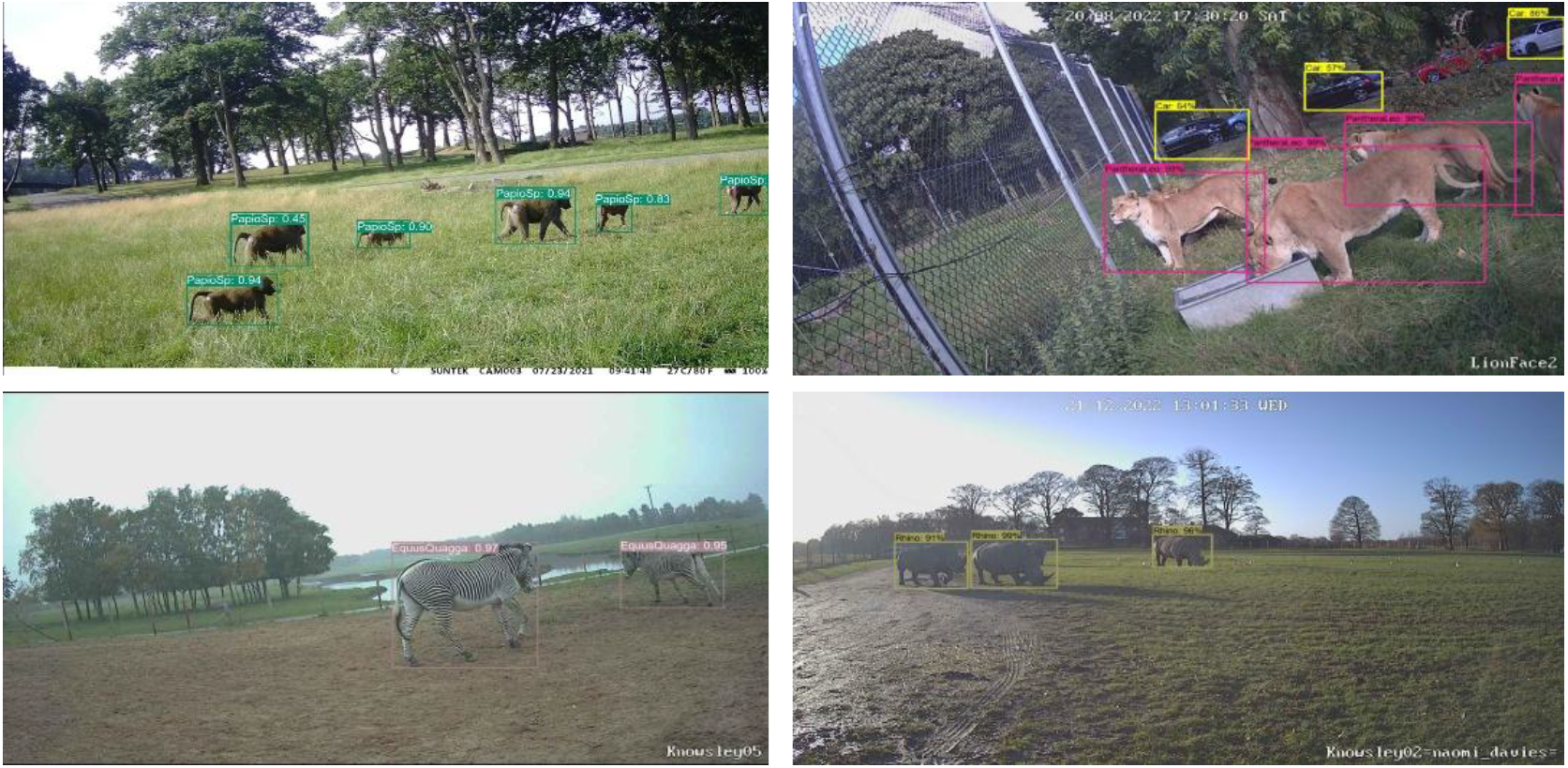
Object detection examples of baboon (top left), lion (top right), zebra (bottom left) and rhino (bottom right). In each of the baboon (CAI=6, HUM=6), zebra (CAI=2, HUM=2) and rhino (CAI=3, HUM=3) images there is agreement between CAI and human observer. In the lion image there is an example of a difference in detection rate (CAI=5, HUM=5).

All statistical analyses were conducted using R 4.3.3. To assess the difference between the two observation methods, a paired Wilcoxon signed-rank test was performed for each species due to the data not meeting the assumptions of normality. This test compared the difference in the number of animals seen across each image for each species. The test statistic (V) was calculated, and p-values were reported to determine the significance of the results. Following this, a Spearman’s rank correlation coefficient was used to test the strength of the relationship between each methodology.

### 2.5 Ethics Statement

This research was reviewed and approved by the Liverpool John Moores University Research Ethics Committee under reference number SW_NJW/2022-5. The study adhered to all relevant institutional and national guidelines for research involving animals and conforms to UK legislation under the Animals (Scientific Procedures) Act (1986).

## 3. Results

### 3.1 Time comparison

When timed, the trained human observer conducted count assessments of 3015 images of baboons in 4 hours, 52 minutes. In direct comparison, CAI classified the same set of images in 52 minutes. This equates to an 82% reduction in processing time.

### 3.2 Classification performance

The performance of the Sub-Saharan Africa model was evaluated using four confusion matrices, one per species included in this study. Lions presented a high number of true positives and true negatives (TP=1224, TN=6058) along with relatively low false positives and false negatives (FP=343, FN=115). The model performed best when classifying rhino, which presented a high number of true positives and true negatives (TP=2629, TN=5016) along with very low false positives and false negatives (FP=6, FN=28). Zebra presented a high number of true positives and true negatives (TP=1349, TN=5680) along with relatively low false positives and false negatives (FP=115, FN=27). Baboons presented a high number of true positives and true negatives (TP=1924, TN=6181) along with relatively low false positives and false negatives (FP=41, FN=139). These results show that for each species included in this study, the Sub-Saharan Africa model performs to a high standard, with the above figures producing high accuracy and F1 scores across the board (Table 1).

**Table 1.**
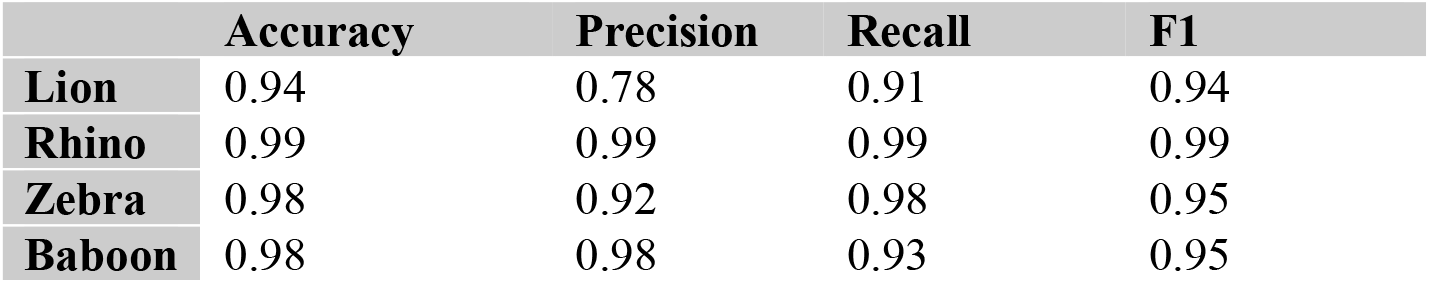
Confusion matrix results for each of the four species studied.

|  | Accuracy | Precision | Recall | F1 |
| --- | --- | --- | --- | --- |
| **Lion** | 0.94 | 0.78 | 0.91 | 0.94 |
| **Rhino** | 0.99 | 0.99 | 0.99 | 0.99 |
| **Zebra** | 0.98 | 0.92 | 0.98 | 0.95 |
| **Baboon** | 0.98 | 0.98 | 0.93 | 0.95 |

### 3.3 Count comparison: Conservation AI compared to a trained human observer

Overall counts of individuals conducted by both human observer and CAI across images in each species show the highest degree of similarity in zebra (CAI = 380, HUM = 379), followed by lion (CAI = 671, HUM =683) and rhino (CAI = 578, HUM=622), with the most disparity in number of individuals observed seen in baboons (CAI = 2006, HUM = 2528).

The Wilcoxon signed-rank tests showed significant differences in the number of individuals observed in images by a trained human observer and CAI, in lions (V=116, p<0.005), rhino (V=310, p<0.005), and baboon (V=128, p<0.005). It showed a non-significant difference between a trained human observer and CAI in zebra (V=105.5, p>0.05).

Results of the Spearmans rank correlation test showed that there was a strong positive correlation between human observer and CAI when looking at the number of individuals in images of each of the four species studied; baboon (rho=0.955, p<0.005), lion (rho = 0.969, p<0.005), rhino (rho=0.887, p<0.005) and zebra (rho=0.843, p<0.005) (Figure 3).

**Figure 3.**
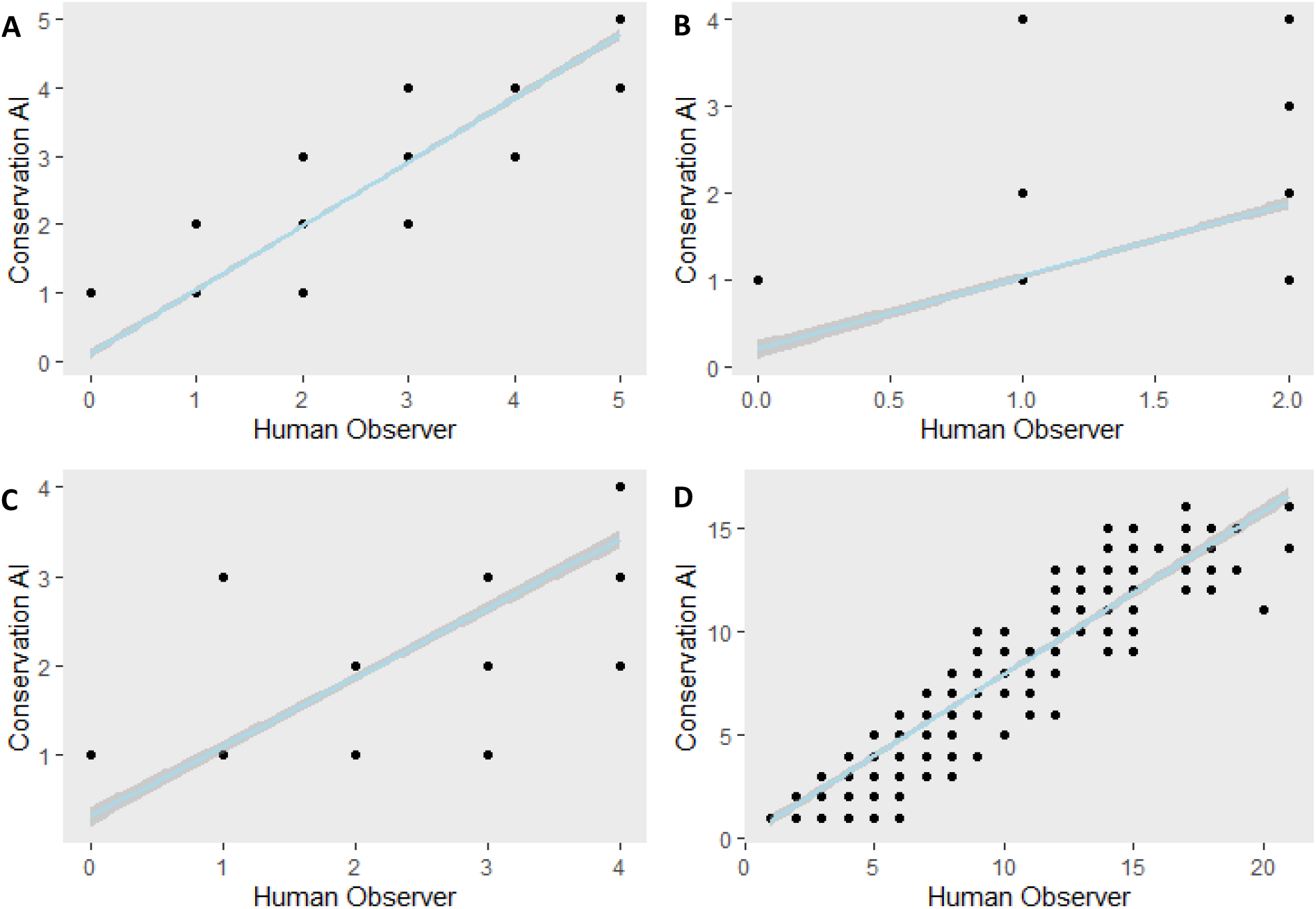
Scatter plots demonstrating the positive correlation between CAI observations vs. a trained human observer in four species; lion (A), zebra (B), rhino (C) and baboon (D).

## 4. Discussion

### 4.1 Findings

This research shows that image recognition can be an effective tool in monitoring select zoo-housed species. This is especially true when balancing error margins against the high cost of time and effort for zoo staff in traditional monitoring. The high accuracy rate highlighted for each species in the confusion matrices, and strong correlations between human and CAI observation shown in each of the species trialed, also allow for confidence to continue development for further use in a zoo environment for monitoring of animal behavior and welfare. The time difference shown between the two methodologies on display is a positive time saver for zoo staff in itself.

However, when considering the passive nature of classification facilitated by CAI, versus an arguably repetitive and time-consuming task, it opens conversations about the costs and benefits of each method. Although this study represents an initial stride forward and shows high levels of accuracy in species identification, it must be noted that count disparities produced significant differences between the trained human observer and AI in three of the four species included in this study. Notably, for rhino and lion, the overall count difference was minimal. Still, instances of animals overlapping, or being significantly occluded by enclosure features or the edge of the image caused minimal disparities in the count, enough to show a lack of agreement across the dataset during analysis. In these cases, the strong positive correlation between the two observation methods showed overall agreement, allowing us to have confidence in CAI as a methodology when considering the automatic identification of these species within Knowsley Safari.

Where the model performed well in the accuracy of baboon identification, and a strong positive correlation between methodologies was still displayed, higher disparities in count data when compared to the other species included in this study show the need for further development to allow for more accurate counting, especially in more complex environments. The baboon enclosure at Knowsley Safari presents a unique set of difficulties, with a high animal density and occlusion of the species by vegetation due to their relatively small body size. Although we highlight the factors specific to this dataset, there are proven accuracy issues with smaller species and species in more cryptic environments (Delplanque et al., 2023). These accuracy issues are exemplified in modern zoos where naturalistic enclosures and group dynamics are favored to promote high welfare standards (Beer et al., 2023). There are also known weaknesses in machine learning models in herd or grouped species due to occlusion and scale variation (Deplanque et al., 2023). Whilst we demonstrate this issue with baboons, it is an important note for zoos hoping to deploy this technology on other group-living species where sociality is a key component of their life and impacts exhibit development and husbandry techniques (Beisner et al., 2023). The higher count disparity and significant difference between CAI results and the human observer in the baboon data set demonstrates a weakness in the model that we can link specifically to the husbandry and life history of baboons. However, this can also likely be extrapolated beyond baboons. Further training of the models across various species within multiple zoological institutions will allow the model to perform better, and this research will realize an all-encompassing zoo monitoring system.

This research is a pivotal starting point in demonstrating novel ways species recognition can reduce workload for zoo staff. Even at the most basic level, the ability to filter camera trap data for expedited classification, already widely used in conservation practice (Green et al., 2020; Trnovszky et al., 2017), can reduce the time needed for monitoring and mitigate access limitations, such as monitoring of out-of-hours behavior. Beyond data classification, the ability to utilize 4G camera technology to monitor animals in real time, creates the opportunity for zoo staff to respond proactively and in an evidence-based manner to husbandry and health needs of the animals in their care, an opportunity that is vital for achieving high welfare standards (Rose & Riley, 2021). The automated accumulation of simple data, such as animal detections as demonstrated in this paper, collected over an extended duration, can be incredibly insightful for zoos. By monitoring enclosure use through automated camera triggers and image recognition technology, data can help inform husbandry needs. For example, investigating dynamics in mixed species exhibits can help staff gain insight into animal behavior where interspecies conflict, resource consumption, and management complexity are often of concern (Veasey & Hammer, 2010). Furthermore, deviant behavior, highlighted when a baseline of species enclosure use is monitored over time through the simple detection of animal presence, can be used to gain insight into the health of species in that environment (Widowski et al., 1990; Kuster et al., 2020) and prompt investigation allowing staff to intervene where necessary. This also benefits welfare assessments, whereby a more species-specific approach can be taken, looking at a broader range of welfare indicators indicated by animal presence and enclosure use based on an individual species’ husbandry requirements.

Taking the heavy resource element away from zoos means they can be more creative with the research possible. Having a baseline monitoring program means the data can be used or added to where needed to facilitate a wider variety of research. This baseline monitoring also helps address some of the common limitations in zoo environments, such as multiple observer studies which can lead to inconsistencies in data reliability and recording (Maher et al., 2021; Brouwers et al., 2021), and the smaller sample sizes that often lead to statistical challenges, affecting publication capacity (Kuhar, 2006; BIAZA Office, 2019). As demonstrated, increased technology integration in zoo research is crucial in extending the scope of zoo research. It is an area that zoos are notably lacking when compared to animal monitoring in situ, or when comparing the use of welfare monitoring technologies to industries like farming (Diana et al., 2021; Madhavan et al., 2021). Developing AI models in zoo environments can also solve models’ lack of training data, leading to misclassification in wild environments (Wearn et al., 2019). This can ensure data availability for a wide range of species to further expand the capacity for automation of animal detections in wild environments, further demonstrating the capacity for zoos to contribute to in-situ conservation efforts beyond the species in their immediate care.

### 4.2 Limitations & counterarguments

Limitations shown in this study’s results can largely be attributed to the inconsistency in data, which is an inherent issue with camera trap data. There is plenty of literature that shows the capabilities of AI to improve when optimally trained (Mohri et al., 2012; Russell & Norivg, 2021; Fergus & Chalmers, 2022), and when considering the variation between zoos, even when housing the same species, it shows the need for a robust approach to training and development. To implement this technology in zoos, substantial effort would be required to develop species-specific models, involving collaboration between developers and zoo staff. Inconsistency of data quality and quantity is largely mitigated by the captive nature of the animals studied. However, issues may arise with more cryptic species and differences in the individual setups and social dynamics of individual species across multiple collections. These factors will make it difficult to standardize a protocol for deployment. We must also consider disparity in access to monitoring technologies across zoological collections. However, there are options for integrating automated monitoring solutions with any setup, from basic camera traps to state-of-the-art systems, ensuring accessibility and adaptability for all institutions. All limitations can be lessened by an interdisciplinary approach to developing these technologies, engaging zoo staff throughout the process to ensure that the models are functional and useful.

### 4.3 Future research

Going forward, the aim is to develop this methodology further towards deploying an all-encompassing zoo monitoring system. The next steps will include standardizing the installation protocol for the most effective monitoring and integrating pose estimation to investigate the potential for automating the study of specific behaviors within a zoo setting. The ability to not only classify animals to a species level, but also incorporate behavioral analysis widens the scope for zoo research, extending the possibilities for longer-term multi-zoo studies and moving further towards the automation of behavior assessments. As AI technology advances, expanding its capability to provide richer, multifaceted data on individual animal behavior, health, and sociality will be an important direction for future research to provide more comprehensive insights beyond species presence.

### 4.4 Conclusion

This research demonstrates that image recognition technology can effectively monitor zoo-housed species, providing significant time savings and high accuracy in species identification compared to traditional methods. However, challenges remain in accurately counting animals in complex environments, particularly for smaller and high-density species like baboons, where occlusion and group dynamics pose difficulties. These limitations highlight the need for further model development to enhance performance in differing conditions, something that is vital for universal use across zoos. Despite these challenges, the technology substantially benefits zoo staff, allowing for reduced workload and more proactive, evidence-based animal welfare management. Future research should focus on refining species recognition, standardizing installation protocols across zoos, and integrating behavioral analysis into the monitoring process. This will expand the technology’s capability to provide a more comprehensive and automated zoo monitoring system, enabling longer-term, multi-zoo studies that improve both animal behavior and welfare assessment.

## Acknowledgements

The authors would like to thank Vodafone for their donation of SIM cards to the wider project, a proportion of which were utilized to collect data for this study. We would also like to thank the staff at Knowsley Safari for their ongoing support of this research, specifically Chris Smart for organising camera access, and finally Jonathan Cracknell for his contribution to the preparation of a figure for the manuscript.

## Conflict of Interest Disclosure

The authors declare no conflict of interest that could influence the work reported in this paper.

## Data Availability Statement

The data supporting the findings of this study are available from the corresponding author upon request.

## Notes

### Competing Interest Statement

The authors have declared no competing interest.

